# EEG in game user analysis: A framework for expertise classification during gameplay

**DOI:** 10.1101/2021.01.29.428766

**Authors:** Tehmina Hafeez, Sanay Muhammad Umar Saeed, Aamir Arsalan, Syed Muhammad Anwar, Muhammad Usman Ashraf, Khalid Alsubhi

## Abstract

Video games have become a ubiquitous part of demographically diverse cultures. Numerous studies have focused on analyzing the cognitive aspects involved in game playing that could help provide an optimal gaming experience level by improving video game design. To this end, we present a framework for classifying the game player’s expertise level using wearable electroencephalography (EEG) headset. We hypothesize that expert/novice players’ brain activity is different, which can be classified using the frequency domain features extracted from EEG signals of the game player. A systematic channel reduction approach is presented using a correlation-based attribute evaluation method. This approach identifies two significant EEG channels, i.e., AF3 and P7, from the Emotiv EPOC headset’s fourteen channels. The features extracted from these EEG channels contribute the most to the video game player’s expertise level classification. This finding is validated by performing statistical analysis (t-test) over the extracted features. Moreover, among multiple classifiers used, K-nearest neighbor is the best classifier in classifying the game player’s expertise level with up to 98.04% classification accuracy.

**Author summary:** Tehmina Hafeez ROLES Investigation, Writing – original draft * AFFILIATION Department of Computer Engineering, University of Engineering and Technology, Taxila, 47050, Pakistan.

Sanay Muhammad Umar Saeed (Corresponding author) ROLES Conceptualization, Writing – review editing * AFFILIATION Department of Computer Engineering, University of Engineering and Technology, Taxila, 47050, Pakistan.

Aamir Arsalan ROLES Methodology, Writing – review editing * AFFILIATION Department of Computer Engineering, University of Engineering and Technology, Taxila, 47050, Pakistan.

Syed Muhammad Anwar ROLES Validation, Writing – review editing * AFFILIATION Department of Software Engineering, University of Engineering and Technology, Taxila, 47050, Pakistan.

Muhammad Usman Ashraf (Corresponding author) ROLES Validation, Writing – review editing * AFFILIATION Department of Computer Science, University of management and Technology, Lahore (Sialkot), 51040, Pakistan.

Khalid Alsubhi ROLES Conceptualization, Writing – review editing AFFILIATION Department of Computer Science, King Abdul Aziz University, Jeddah, Saudi Arabia.

## Introduction

Technology advancements have revolutionized the gaming industry leading to a dynamic shift from computer games to mobile games. This shift has increased the number of active mobile game players to around 2.1 billion, making the gaming industry one of the highest profit earning entertainment industries. In a significant milestone, the video game industry contributed 80% of the $36 billion profit earned from all software-related industries during 2018 in the United States alone [1]. According to a global games market report, it is expected that the gaming market segment will be worth $180.1 billion by the year 2021 [2]. This rapid growth in the gaming industry creates significant competition for game developers to capture the diverse population of video game players. A major factor in game development deals with improving the user experience, which demands an accurate cognitive assessment of game players to improve user interaction and enhance the gaming experience [3, 4].

The term *player experience*, in the context of game user research (GUR), is derived from the discipline of human-computer interaction. In particular, the player experience is formally explored using constructs such as flow, immersion, challenge-skill balance, affect, presence, motivation, and tension [5]. The player experience is commonly measured by self-reported questionnaires [6], which includes immersive experience questionnaire (IEQ) [7], game experience questionnaire (GEQ), in-game GEQ (iGEQ) [8], and game engagement questionnaire (GEnQ) [9, 10]. There are a large number of questionnaires currently in use, yet the convergence in their usability is minimal. This introduces several challenges both in terms of selecting suitable questionnaires and comparing results across different studies. A hybrid approach can be a possible solution to some of the known challenges [11].

Among various questionnaires used for player experience, none of them are aimed towards identifying the expertise level of the game player. Subjective evaluation using self-labeling is widely used for this purpose, which are gradually replaced by objective evaluation methods. Objective evaluation methods perform a comprehensive analysis of the physiological observations recorded from game players during the period of gameplay. Such observations are efficient in analyzing the cognitive and behavioral aspects of the game player. Since the last decade, the cognitive and behavioral analysis of the game player is becoming an emerging research topic in the GUR. It intrigues the researchers and game developers to explore the prime factors that attract a wide audience of game players [12–14].

Physiological measures electrocardiography (ECG) [15], electromyography (EMG) [16], electrodermal activity (EDA) [15, 17], and electroencephalography (EEG), are significantly gaining importance in GUR [18–21]. In particular, games that provide adaptive features in response to brain signals are known as *neurofeedback games*.The usability of physiological measures in the game design of neurofeedback games has been demonstrated using well-designed experiments [22].

Neuro-feedback games related to EEG can be grouped into three categories, active (for instance, brain arena, brain runner, and shooter games), reactive (for instance, checker game, memory game, and space invaders), and passive (for instance, matrix game, brain ball, and mind ninja games) [23]. Contrary to the conventional game control, the neuro-feedback games use brain signals to adapt the game environment, i.e., either control the game speed or adapt the game design according to cognitive and motor skills of the game player [24]. EEG-based neurofeedback is increasingly used for serious games. MindLight is a serious neuro-feedback game that is developed to reduce anxiety in children [25]. Similarly, 2D and 3D EEG-based games such as brain chi, dancing robot, pipe, and escape are successfully implemented for concentration training purposes [26]. Recently a multiplayer car racing game was proposed to improve the attention level of a user [27]. In particular, it was shown that varying game difficulty in correspondence with the variation in the emotions of the game player helps in preserving their interest and maintains player engagement. It is further reported that both healthy and physically impaired people find adaptive neurofeedback games as motivating, attractive, and more challenging than non-adaptive games [13]. Furthermore, such responsive games are found to be successful for the enhancement of attention and cognition in the game players [28].

Herein we propose a method for the classification of the expertise level of the game player. Our proposed framework would use features extracted from the EEG signals observed during the gameplay. The proposed EEG-based game player expertise classification framework can be augmented with an adaptive neurofeedback game design. It will allow adapting the game difficulty according to the variations in the brain activity of the game player. Thus, it will help in maintaining the skill-competence, attention, and cognition of the game player. This is a step towards improving the player experience by assimilating the concepts of dynamic difficulty adjustment (DDA) and neurofeedback in game design.

### Related Work

Electrophysiological methods provide an objective, continuous, and impartial measure for analyzing the player experience [29].EEG has significantly gained importance in GUR and is shown to be an effective tool in the game studies [20]. For instance, EEG was found to be useful in differentiating the stressed and relaxed states of the game player during the racing and first-person shooter games [21]. Moreover, the level of motivation and relaxation in the game players was investigated using the EEG data [29, 30]. Attention differences among video game players and non-video game players were also investigated using EEG signals; by recording the steady-state visual evoked potential and event-related potentials, [31]. The correlation of the EEG frequency bands with valence, arousal, and engagement was investigated to find the best classifier for various cognitive-affective states of the game player [32]. The wearable EEG devices are found to be a discreet, reliable, and portable way for observing and discriminating the cognitive-affective states of the game player [33]. In particular, the Emotiv EPOC headset is a low-cost and widely used device in the game studies .It was reported that Emotiv EPOC could be a better substitute than laboratory-based EEG systems for analyzing the game player experience [34].

In literature, the classification of game player expertise level was performed using EEG and non-EEG based methods [35]. [36] found that support vector machine (SVM) classifier can be an accurate classifier for real-time classification of the game player experience among other machine learning-based classifiers. Real-time analysis of game player experience helps keep the player positively influenced, motivated, immersed, engaged, relaxed, and improve the cognitive abilities [37–39]. While [40], found that the player’s expertise and competency can be better classified using features from the EEG signals and regression-based ridge estimator classifier. In [41, 42], mobile game players’ expertise was classified using EEG signals and machine learning algorithms, where the Naive Bayes (NB) was found to be the best among other classifiers used. While expertise level classification is important, there are limited studies available in the literature [40, 41, 43]. We argue that for GUR, this area is significant and hence needs a more systematic analysis. It is observed that the reported classification accuracy needs to be improved further to develop a better game design based on the expertise level of the game player. These studies also presented a limited analysis of the brain activities concerning the player groups’ expertise level. Herein, we aim to achieve an improved and accurate classification of the expertise level (expert/novice) of the game player with a lesser computational cost. It could be a step towards the development of adaptive neurofeedback games that would adjust the game difficulty in real-time. We hypothesize that the brain activity of expert/novice players is different, which can be classified using the frequency domain features extracted from the EEG signals of the game player.

### Our Contributions

We aim to classify the game player’s expertise level in one of the two categories, i.e., expert or novice, by utilizing features of the most relevant EEG channels. Hence the computational overhead for analysis and real-time implementation of such a system will reduce. Further, we intend to improve the classification accuracy of the game player’s expertise level. Towards this, we used well-established features classifiers but we intend to find the best suited EEG electrodes and features for the task at hand using a wearable device. Henceforth, this study aims to contribute in the following ways in GUR,

1. An improvement in the classification accuracy of the game’s expertise level player during gameplay using features extracted from the EEG signals.
2. A computationally efficient mechanism for the classification of game player expertise level by effective EEG channel selection and feature-length reduction.

The rest of the paper is organized as follows. pm describes the proposed methodology along with the materials and methods used in the study, er presents the experimental results followed by the discussion and conclusion in disc and conc, respectively.

## Proposed Methodology

A block diagram of the proposed scheme for EEG-based game player expertise classification is presented in Fig 1. It consists of three main stages, namely data processing: which includes pre-processing and channel selection, feature extraction, and expertise classification. The EEG data used in this study were collected in a previous study [42] and is available at http://tiny.cc/uiae9y. However, a brief overview of the EEG data and experimental procedure for EEG data acquisition is described here for completeness. The following subsections present details for each of the blocks used in the proposed framework.

**Fig 1.**
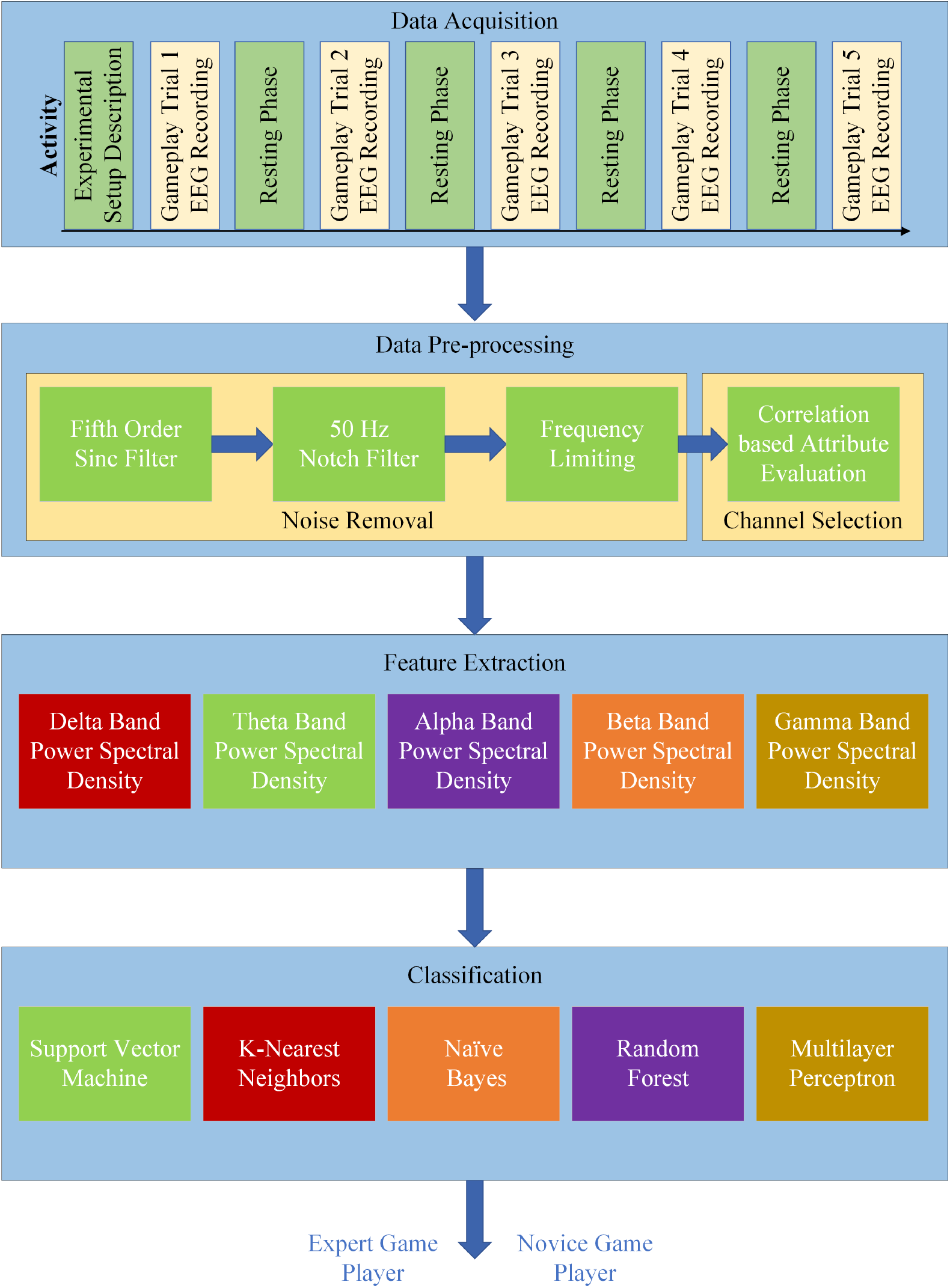
Block diagram of the proposed methodology for the expert-novice classification of the game player.

### EEG Dataset

Ten healthy subjects (one female and nine males), with an average age of 20.37 years, voluntarily participated in the experimental study. All participants belonged to the Asia-Pacific region, having the same educational background, and reported experience in playing games. A popular android based game, *Temple Run*, was used as a stimulus in this experimental study for EEG data acquisition. It is an endless game, which continues until either a demonic monkey eats the character or the character falls in the river or is hit by an obstacle. The experimental protocol followed was designed following the recommendations of the Helsinki declaration.

EEG data acquisition was performed using a 14-channel Emotiv EPOC headset by placing it over the participant’s scalp. Emotiv EPOC is a flexible, high resolution, multi-channel wireless headset and provides fourteen electrodes arranged according to the 10-20 electrode placing system (as shown in Fig 2). These electrodes include AF3, AF4, F7, F8, F3, F4, FC5, FC6, T7, T8, O1, O2, P7, and P8. Further, two additional reference electrodes (CMS and DRL) are positioned on the rear of both ears. The headset provides good temporal resolution and electrodes sufficiently cover the scalp to capture the brain’s electrical activity. For instance, electrodes AF3 and AF4 cover the pre-frontal region, F3, F4, F7, and F8 cover the frontal region, FC5 and FC6 cover the frontal-central region, T7 and T8 cover the temporal region, P7 and P8 cover the parietal region, and O1 and O2 capture the neural activity from the occipital region of the brain. While the signal quality from such wearable devices could be questionable for clinical studies, it has been shown that the acquired data yields significant results in less critical applications.

**Fig 2.**
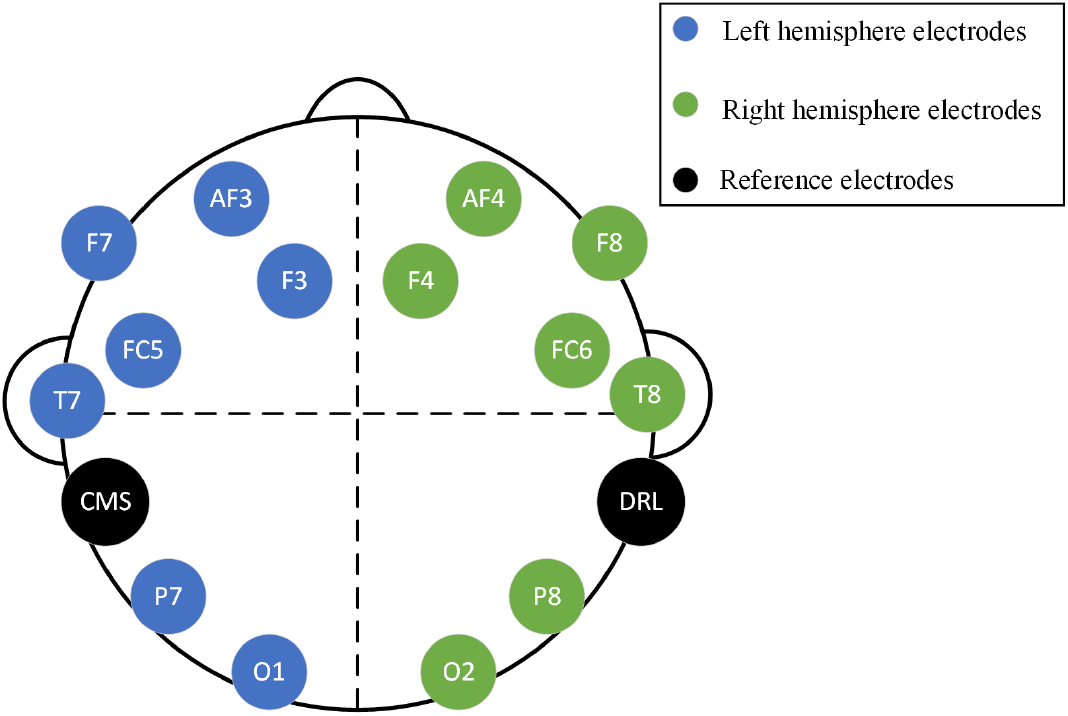
10-20 electrode placing system for placing the Emotiv EPOC headset [42].

During the experiment, EEG electrodes were soaked with a saline liquid to acquire good quality signals. The study was approved by the board of postgraduate studies, Department of Computer engineering, UET Taxila. All participants were informed about the experimental procedure, and written consent was obtained. It was followed by filling the demographic sheet comprising information including name, age, education, and gender. The participants were provided with a comfortable chair and a smartphone with a pre-installed Temple Run game. The room chosen for EEG data acquisition was acoustically noise-free. Moreover, the electric cabling was avoided near the experimental setup to keep the environmental noise to a bare minimum. Additionally, proper synchronization was done between the Emotiv EPOC headset and Emotiv software development kit (SDK) to avoid erroneous or faulty data acquisition.

EEG data of each participant was recorded for five trials of gameplay. Each trial of the gameplay was separated by a resting phase of 60 seconds. The average gameplay time observed for each participant was 24.7 minutes. EEG data were recorded at a sampling rate of 128Hz and was saved in the European data format (EDF) using Emotiv SDK. The recorded data was transferred via Bluetooth to a computer for further offline analysis. Game score after each trial of the participant was recorded, and an average score was calculated for all of the participant’s five trials. A threshold (*T*) on the game score of each participant was calculated by adding the scores of all trials of all participants and dividing it by the total number of participants [41]. The mathematical representation of the threshold (*T*) is calculated as follows.

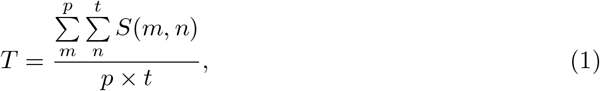

where *p* is the total number of participants, *t* is the total number of trials, and *S(m,n)* is the score of *n^th^* trial of the *m^th^* participant. The participant with an average game score above this threshold was labeled as an expert while lower than this was labeled as a novice. These assigned labels also corresponded to the subject’s self-reported expertise. Thus, four participants were identified as an expert, while six participants were identified as novice based on the threshold value.

### Data Processing

The data processing stage consists of two sub-stages, i.e., pre-processing for noise removal and channel selection phase for the selection of the most significant EEG channels. The output of this stage, i.e., data processing, identifies relevant EEG channels whose features contribute most in classifying game player expertise level. The details are presented in the following subsections.

### Preprocessing

EEG signals have micro-volt amplitude making them highly prone to contamination by various artifacts. These artifacts, including muscle movement, electrical lines, and electrode movement, should be carefully removed prior to the further processing of the EEG data. Artifacts such as eye blinks were automatically detected and recorded by the Emotiv SDK and were removed manually or by frequency limiting procedures [32]. Emotiv EPOC has a built-in 5^*th*^ order sinc filter, which acts as a low pass filter with a low cut off frequency and higher attenuation of stop-band to reduce noise in EEG signals. It also has notch filters at 50 and 60 Hz, which acts as a band-stop filter to rectify noise generated by power supply lines.

### Channel Selection

After the pre-processing step, the EEG signals were subjected to channel selection, where the average power of time-domain EEG signals was computed across all fourteen channels. Mathematically average power was calculated as follows.

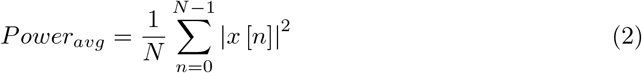

where *x* is the EEG signal with length *N* across each EEG channel. After time-domain power computation, *correlation-based feature selection* was applied to select the most relevant channels for game player’s expertise classification [44].

*Correlation-based attribute selection* is based on the principle that an attribute having a high correlation with the class labels must be selected, whereas attributes which have a high correlation with other attributes should be ignored. The correlation between each attribute and output class was mathematically calculated as follows.

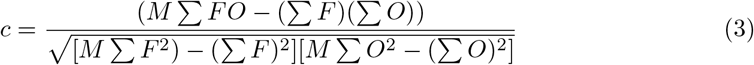

where *M* is the total number of pair of scores, *F* is the feature vector, *O* is the output class, *c* is the *Pearson's correlation coefficient*, ∑*F* is the sum of *F* scores, ∑*FO* is the sum of products of paired scores, ∑*F*^2^ is the squared *F* scores, ∑*O* is the sum of output class and ∑*O*^2^ is the sum of squared *O* scores.

The correlation-based feature selection was applied in *Weka* tool using the *Correlation AttributeEval* method, which evaluates the correlation of all fourteen EEG channels with the output class and presents correlation coefficients in a ranked order. Out of the fourteen EEG channels, the top-ranked channels were chosen, which were highly correlated with the output class to reduce the computational overhead and avoid the chances of over-fitting. Further, the features from these selected channels were extracted for the game player’s expertise classification.

### Feature Extraction

In general, *power spectral density (PSD)* represents the power distribution of a signal over a particular frequency. In this study, the analysis of the recorded EEG data was performed in the frequency domain by computing the PSD of the time domain signal. To this end, *Welch* method was employed using *Brainstorm3* toolbox in MATLAB. The *Welch* method was used to compute the power spectrum of the input signal by dividing it into 50% overlapping sequences of the specified length followed by the application of the *Hamming* window to each of the overlapped sequences. Further, *Fast Fourier Transform (FFT)* of each sub-region was computed. Finally, power from the FFT coefficients of all overlapped windows was averaged. Thus, the time-domain EEG signal was represented by the power spectrum of its frequency bands [45]. The extracted PSD features from each channel included EEG frequency bands in the following ranges: *delta (2-4 Hz)*, *theta (5-7 Hz)*, *alpha (8-12 Hz)*, *beta (13-29 Hz)*, *gamma1 (30-59 Hz)* and *gamma2 (60-90 Hz)*. The default settings of *Brainstorm3* resulted in *gamma* values in two different ranges i.e., lower gamma as *gamma*_1_, and upper gamma as *gamma*_2_. These features were extracted for those EEG channels which resulted in a high correlation with the output class and hence were selected as discussed in channelselection.

### Game Player Expertise Classification

For the expert-novice classification problem of the game player, five supervised machine learning algorithms were used, including support vector machine (SVM), the Naïve Bayes (NB), K-nearest neighbor (KNN), multiplayer perceptron (MLP), and random forest (RF). The details of these algorithms are presented in the following text for completeness.

#### Support Vector Machine (SVM)

SVM is a supervised machine learning algorithm used for both classification and regression problems [46]. It uses linear, non-linear, Gaussian, or polynomial kernels to achieve a hyperplane that efficiently separates the input data. Herein, we classified the input feature set in two classes, i.e., expert and novice, using the output class labels and used a linear kernel. In a previous study [42], game player expertise was classified with 80% accuracy using an SVM classifier. In another study, SVM had classified the game player’s expertise with 82% and 86% classification accuracy when features of 14 and 4 EEG channels were utilized respectively [41]. Apart from game player expertise classification, SVM is well suited for the classification of emotions in general gameplay events [32]. As observed in the previous studies, the efficiency of the SVM algorithm provided a rationale for using it in the current study for classifying the expertise level of a game player.

#### Naive Bayes (NB)

NB is a simple statistical algorithm based on the Bayes Theorem. It is based on the class conditional independence assumption, which states that the effect of the predictor on the given class is independent of the values of other predictor classes [47]. This assumption makes the algorithm robust and computationally cost-effective to be used in real-time applications. Despite the fact that NB is a simple statistical algorithm, its usability in this study is for the reason that NB does not require a large training data set for the accurate classification [32]. In the context of game player expertise classification, NB is found to be an effective classifier with the 88.89% classification accuracy [42]. In another study, NB classified the expertise of the game player with 84% and 88% classification accuracy when features from 4 and 14 EEG channels were used, respectively [41].

#### K-Nearest Neighbor (KNN)

KNN is one of the simplest supervised machine learning algorithms widely used in varied problems because of its low computational complexity. The classification process of KNN is relatively simpler as it analyses the feature similarity of each input data point with the output class and classifies the input data point to the output class with which it has the maximum correspondence. It classifies the input data point by finding the distance or closeness of its features from each of the output classes, i.e., expert or novice. The K value in KNN denotes the number of nearest neighbors. The value of *K* chosen in this study is 1. In the context of classifying the game player emotions, KNN is found to be a good classifier. A study conducted by Lin et al. [48] has verified the usability of the KNN classifier in classifying the player experience. Their study’s results have reported that the KNN classifies the four emotions with an accuracy of 82% using the features of the EEG signals. In another study, five different emotions were classified with an accuracy of 82.87%, and 78.57% for the data obtained from 62 and 24 EEG channels when the stimulus used were emotional audio-visual clips [49]. The study conducted by Parsons et al. has identified that the KNN is the best classifier for classifying the general game based events when the beta band is used as a predictor feature [32]. The previous studies in the GUR have used the KNN classifier to classify various player experiences, whereas none has utilized it for the classification of the game player expertise. Nonetheless, the efficiency of the KNN classifier is well known, and because of its optimum performance in the previous studies, it is chosen in the current study for the classification of the game player expertise using the EEG features.

#### Multilayer Perceptron (MLP)

MLP is a type of neural network that uses the backpropagation algorithm for classification purposes. This feed-forward neural network-based algorithm works by assigning weights to the input data, which are mapped to each neuron’s output through the transfer function. Thus, a non-linear relationship is established between weighted inputs to the output of the network. The frequently used transfer functions in the MLP classifier are sigmoid, hyperbolic tangent, rectified linear unit, and radial [50]. The transfer function used in this study is the *sigmoid function* and the hidden layers are chosen in *Weka* as parameter *a* i.e. the number of attributes + number of classes / 2. In the context of game player expertise classification, MLP has proved to be a good classifier with the 80% accuracy of expertise classification [42]. In another study, the MLP classifier achieved 82% and 84% classification accuracy when features from 14 and 4 EEG channels were used, respectively [41]. The performance of MLP in the previous studies endorses its use in the current study and future studies.

#### Random Forest (RF)

RF is a supervised learning classifier that is widely used because of its robustness, insensitivity to overfitting, and ability to model the categorical values. Its applicability is suited for classification as well as regression tasks. It is an ensemble algorithm that works by creating multiple decision trees by randomly selecting features from the training set and then predicting the test data class by finding out its similarity to all of the decision trees in the forest. RF is an effective classifier for the classification of multiple user states based on their EEG signals while the users are engaged in the game playing [51]. By reviewing the literature, it has been identified that RF is neither utilized for the game player expertise classification nor the classification of player experience. Nonetheless, the applicability and usability of the RF classifier is well known, and therefore, it is used in the current study for the EEG based game player expertise classification.

#### Classification-Validation Model

For every classifier, the performance of classification is assessed using various evaluation parameters. In this study, the performance of the various classifiers was evaluated using 10-fold cross-validation (CV) model where all of the trial instances of the game players were divided into 10 splits; 9 splits were used for training the classifier and 1 split for testing the classifier performance. We further used leave-one-out cross validation (LOOCV) and leave-one-subject-out (LOSO) cross validation for evaluating our models. Since the data used in this study was limited, LOOCV was used to verify the classification results along with the 10-fold CV. While LOSO was used to analyze the results for subject-based analysis instead of using all instances independently.

#### Performance Evaluation Parameters of Classification

The performance metrics selected in this study for analyzing the classifier performance are well known and used in other studies as well ([42] and [41]). The classifier performance of the various classifiers was evaluated in terms of classification accuracy and error performance parameters. Other than classification accuracy, performance parameters such as precision, recall, F-measure, and receiver operating characteristic curve (ROC) were also evaluated for each classifier. Since the data used has an imbalance in class labels, the evaluation parameters, i.e., precision, recall, and ROC analysis, were included to verify the accuracy. Furthermore, every classifier was also evaluated for its error performance using mean absolute error (MAE), root mean square error (RMSE), relative absolute error (RAE), and root relative squared error (RRSE). The performance measures were calculated as follows.

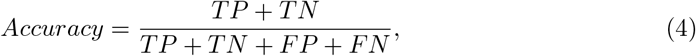

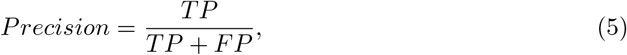

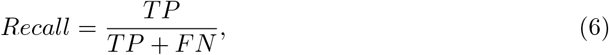

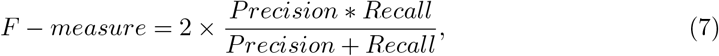

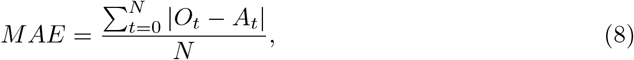

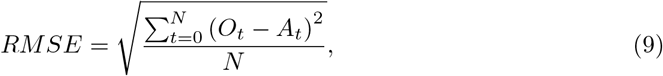

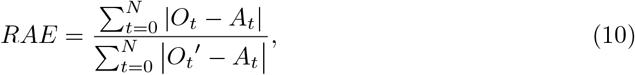

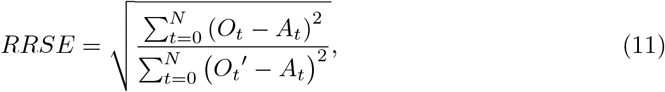

where *N* depicts the total number of observations and *A_t_*, *O_t_* and 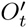 denote actual, observed and mean of all the predicted observations. Furthermore, true positive, true negative, false positive and false negative observations are denoted by *TP*, *TN*, *FP*, *FN* respectively. ROC represents the plot between the true positive rate (TPR) and the false positive rate (FPR) of the classifier.

## Results

The EEG data obtained from the 10 participants were classified into expert and novice categories by thresholding each participant’s average game score (as described in the EEG Dataset subsection) followed by the preprocessing step to remove artifacts. The clean EEG signals were then fed to the channel selection step, where each EEG channel’s power was treated as an attribute of interest. The output of this block gave the EEG channels, which were maximally related to the output class and whose features were used to classify the game player expertise. The results from each of these stages are presented in the following subsections.

### Performance of Channel Selection

Channel selection performed using *correlation-based attribute selection* method evaluated the correlation of the power of each of the fourteen EEG channels with the output class. *Pearson’s Correlation Coefficient* found for all fourteen EEG channels is shown in Fig 3 The highly correlated EEG channels were chosen using the threshold, 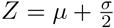, where *μ* and *σ* denote the *mean* and *standard deviation* of the correlation coefficients of the EEG channels. The threshold *Z* gave a value of 0.1366, which suggested that EEG channels O1, AF3, P7, and T7 are highly correlated with the output class compared to the rest of the EEG channels. Channel O1 shows the highest correlation with the output class and has a correlation coefficient value of 0.2408. While AF3, P7, and T7 are ranked in second, third, and fourth-order with correlation coefficient values of 0.1908, 0.1409, and 0.1406 respectively. The rest of the EEG channels, whose correlation value was less than *Z*, were neglected due to their decreasing correlation with the output class. It is clear from Fig 3 that the correlation values of EEG channels P8 and T8 approached zero, indicating the least correlation with the output class.

**Fig 3.**
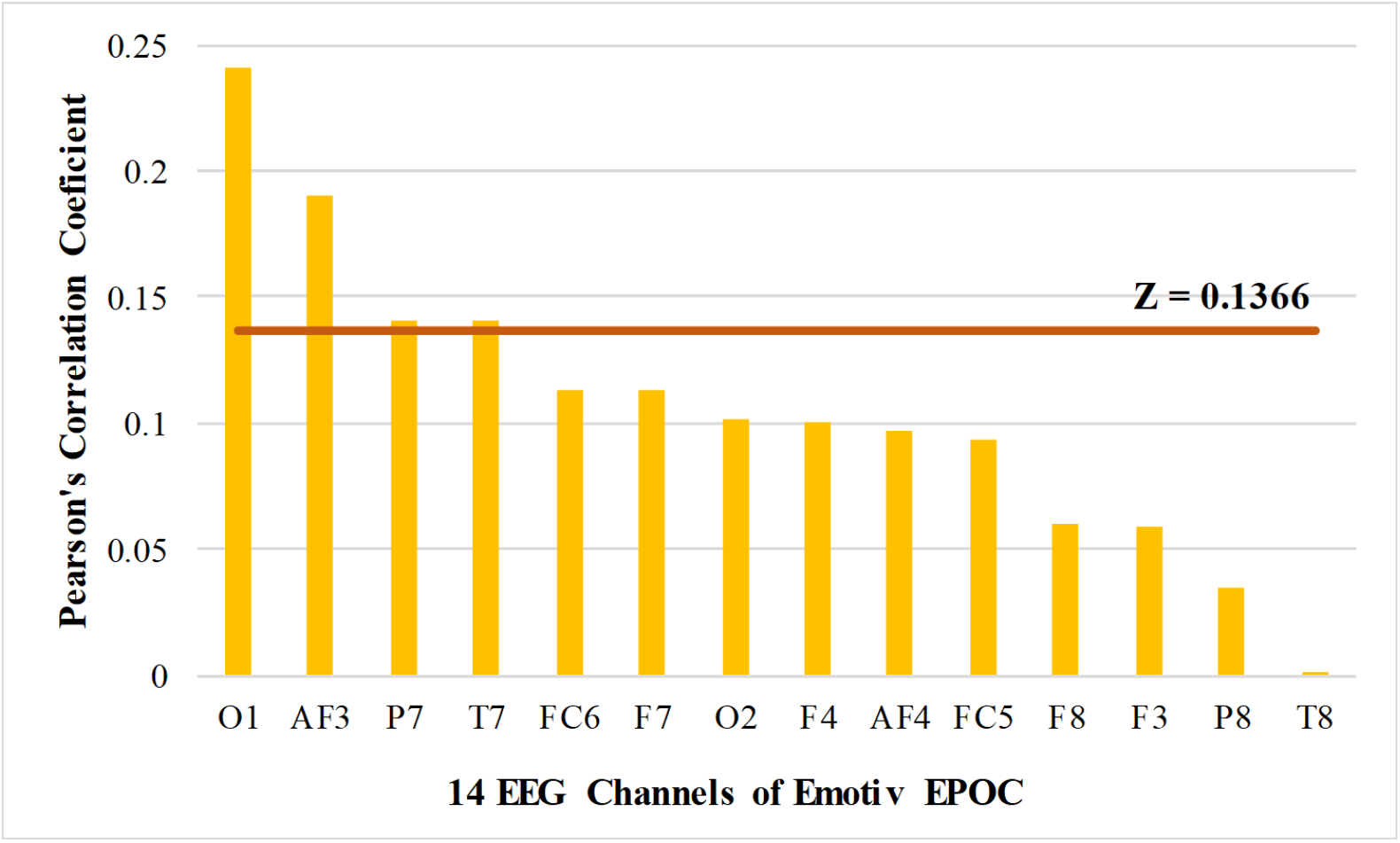
Pearson’s correlation coefficient values of the fourteen channels of Emotiv EPOC obtained by the Correlation-based attribute selection method.

### Classification Performance of the Classifiers

Channel selection resulted in identifying four significant channels, namely, O1, AF3, P7, and T7, whose features were used in the feature extraction phase. PSD features of the four significant channels were computed by using the Welch method resulting in five frequency bands. Thus, for four EEG channels, their corresponding six PSD features were computed and arranged in the form of a matrix. A total of 51 trial files were used, consisting of 19 experts and 32 novice instances, and each channel resulted in six PSD features. Thus, 24 features from the four significant channels were extracted. Thus, a feature matrix of 51 × 24 dimension was formulated accompanied by the class labels for each instance. The rows of the matrix correspond to the total trial instances of the gameplay, and columns correspond to the PSD features. To get a better insight of which channels contribute most in classifying the game player’s expertise, all combinations of the significant EEG channels O1, AF3, P7, and T7 were computed using the combinatorics formula 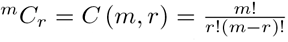 where *m* is the total number of channels, i.e., four and *r* is the number of channels selected from a total of four channels considering no repetitions of channels. This was done to explore which channel or combination of the channel’s features gives maximum accuracy for classifying the expertise of the game player. Thus, fifteen combinations of the EEG channels (4*C*1 + 4*C*2 + 4*C*3 + 4*C*4) are found, and subsequently, all of their features were analyzed for the classification performance. The features of the selected channels were then fed to the classification algorithms to classify the game player’s expertise. Moreover, this classification procedure led to selecting the best classifier that achieved the maximum classification accuracy.

A 10-fold cross-validation model was used for evaluating the classification performance of the features of all fifteen combinations of four EEG channels. Fig 4 depicts the accuracy of game player expertise classification when all fifteen combinations of the significant EEG channels O1, AF3, P7, and T7 were used. The game player’s expertise’s classification results were evaluated for five classifiers SVM, NB, KNN, MLP, and RF. It is visible from Fig 4 that the highest accuracy of 98.04% was achieved through the classification of the KNN classifier using all the features of AF3 and P7 channels. However, using all channels of all channels gives the highest classification accuracy of 96.08%, also achieved through the KNN classifier classification. It is observed that the KNN classifier efficiently classifies the game player expertise using the features of only two significant channels. Our results indicate that using only AF3 and P7 channels gives better classification accuracy compared to other channels or their combination. The improved accuracy of the game player’s expertise classification with less computation achieved by the channel reduction method makes it suitable for the real-time implementation of the DDA based neurofeedback games.

**Fig 4.**
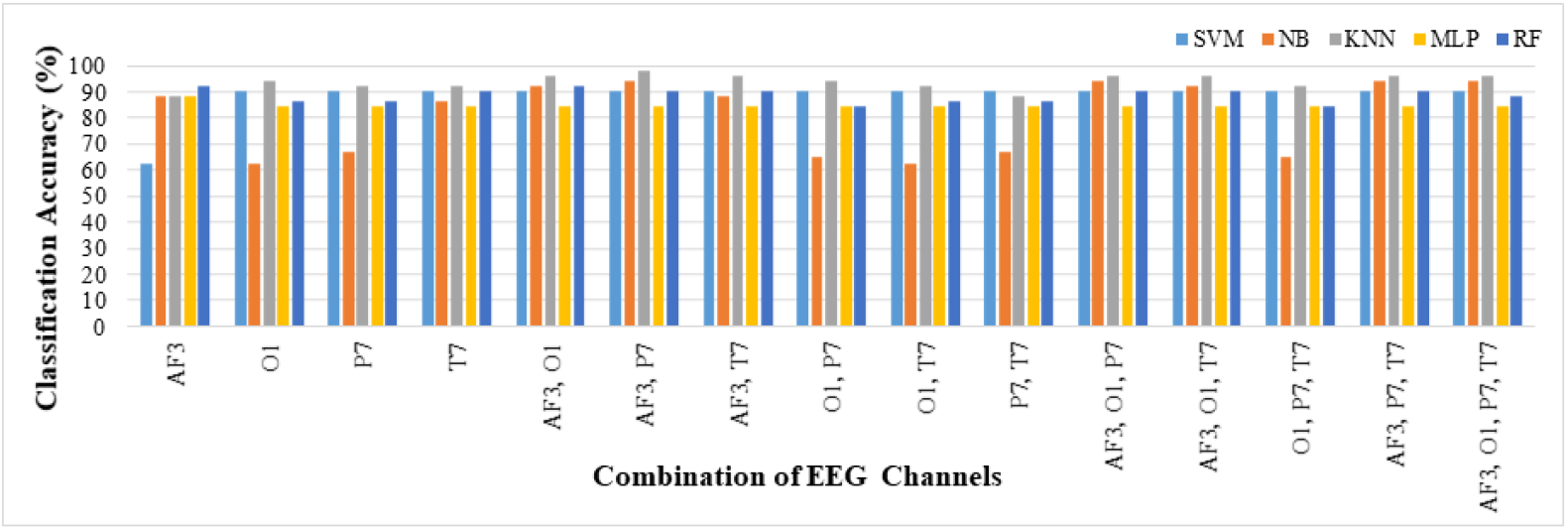
Classification accuracy of the game player’s expertise for fifteen combinations of the EEG channels.

Table 1 depicts the performance metrics for all combinations of the EEG channels that achieve the least accuracy of 96.08% and F-measure of 0.96. These two parameters show the stability of the classification system. It is worth noting that in all cases, KNN is the best classifier for classifying the expertise of the game player based on accuracy and F-measure, which is the harmonic mean of precision and recall. Moreover, AF3 is a significant EEG channel contributing most to the game player’s expertise classification. This active frontal region’s contribution is because game playing involves thought processing, cognition, and decision-making, along with the muscle movements [41, 52]. This thought processing and increased activity of the brain’s frontal region is observed from the significant contribution of the AF3 channel in the classification process. Thus, it is one of the most active regions of the brain that depicts the neural activities that contribute most to classifying the game player’s expertise. The parietal brain region is more related to the processing of spatial information. The active processing of this channel is analyzed by the contribution of its features in the classification process.

**Table 1.** Performance comparison of the KNN classifier for the different combination of the EEG channels used.

| Channels | Accuracy (%) | Precision | Recall | F-measure | ROC |
| --- | --- | --- | --- | --- | --- |
| AF3,P7 | 98.04 | 0.98 | 0.98 | 0.98 | 0.98 |
| AF3,O1 | 96.08 | 0.96 | 0.96 | 0.96 | 0.97 |
| AF3,T7 | 96.08 | 0.96 | 0.96 | 0.96 | 0.95 |
| AF3,O1,P7 | 96.08 | 0.96 | 0.96 | 0.96 | 0.97 |
| AF3,O1,T7 | 96.08 | 0.96 | 0.96 | 0.96 | 0.97 |
| AF3,P7,T7 | 96.08 | 0.96 | 0.96 | 0.96 | 0.95 |
| AF3,O1,P7,T7 | 96.08 | 0.96 | 0.96 | 0.96 | 0.97 |

Fig 5 depicts the error magnitudes for all combinations of the EEG channels that have achieved an accuracy of 96.08% or higher. It is observed that the channel combination of AF3 and P7, whose features have classified the expertise of the game player with 98.04% accuracy, produces the minimum error magnitudes for all error metrics. The error magnitudes of MAE, RMSE, RAE, and RRSE are lowest when features of AF3 and P7 channels are used in conjunction. Henceforth, the features of AF3 and P7 channels not only achieves the highest classification accuracy but also produce the minimum error of classification. Thus, our results show that these two EEG channels are significant for capturing brain recordings, while KNN is the best classifier in such game evaluation scenarios.

**Fig 5.**
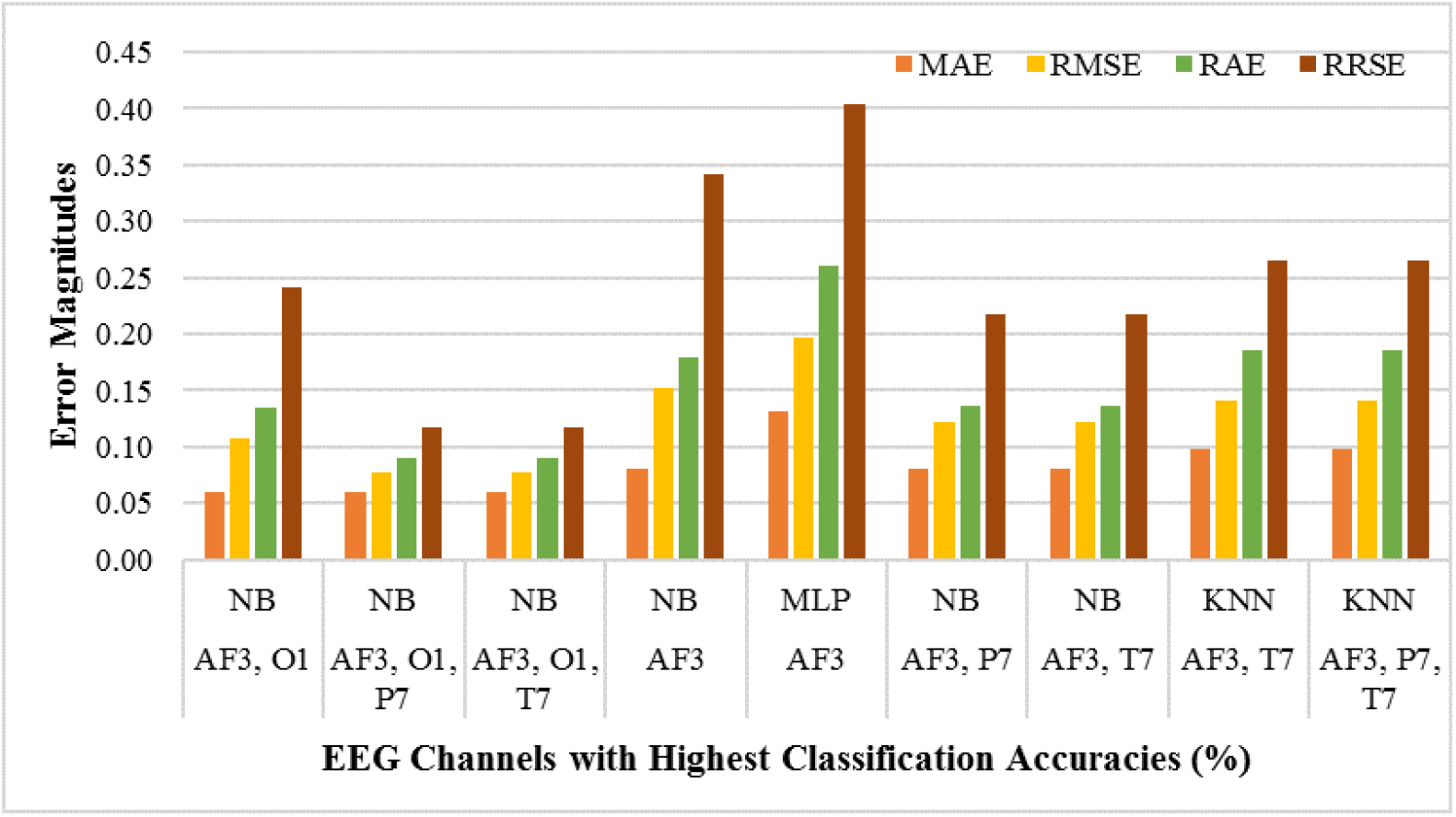
Error magnitudes of the highest accuracy channels classified using K-NN classifier.

### Statistical Analysis

Statistical analysis (t-test) has been performed on the PSD features of AF3 and P7 channels to confirm the obtained results’ validity. A two-tailed t-test confirmed the significance of these features as all of the features have a *p*-value less than 0.05. To get a better insight into the brain activity of the two player groups, the distributional characteristics of the AF3 and P7 channels of the expert and novice player groups were analyzed through box plots that identified the differences in various distributional characteristics frequency bands of the two-player groups. The lower and upper 25% of the data distribution in the box plot represents the lower and upper quartiles, respectively. These are represented by the whiskers, whereas the rest of 50% of the inter-quartile range (IQR) is represented by a box. The horizontal line in the box represents the median of the IQR. The boxplots for PSD features of EEG channel AF3 and P7 are represented by Fig 6 and Fig 7 respectively. The magnitude of PSD is normalized in the range 0 to 1. It is evident that the PSD magnitude of the expert group is visually different from the novice group. Thus, it is obvious from both figures that there is a clear visual distinction for all the EEG bands between the expert and novice groups during the game playing, which verifies our hypothesis that the brain activity of the expert group is different from the novice group and supports the findings of the previous studies [40–42].

**Fig 6.**
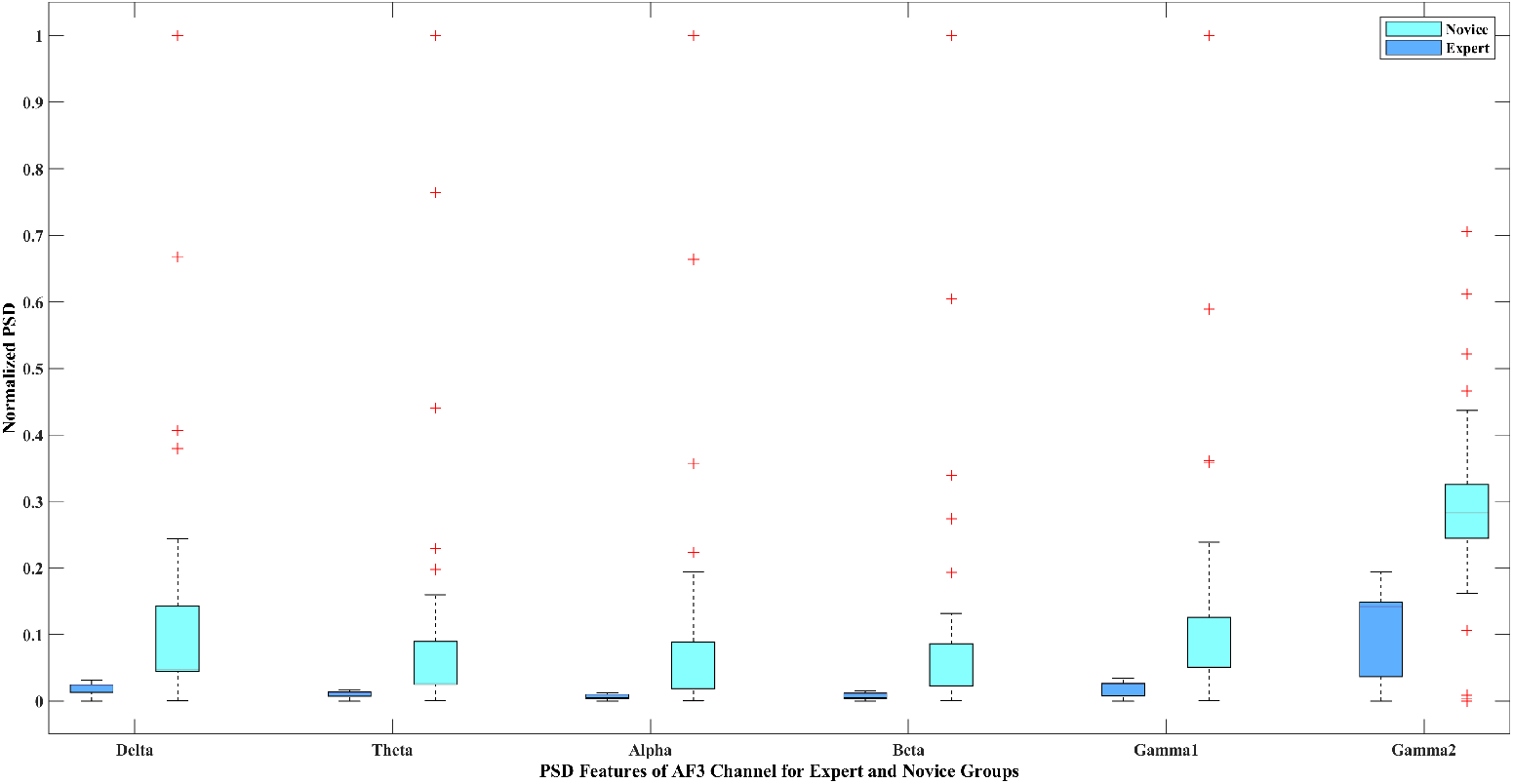
Box-plot representation of the PSD features of *AF*3 channel for expert and novice groups.

**Fig 7.**
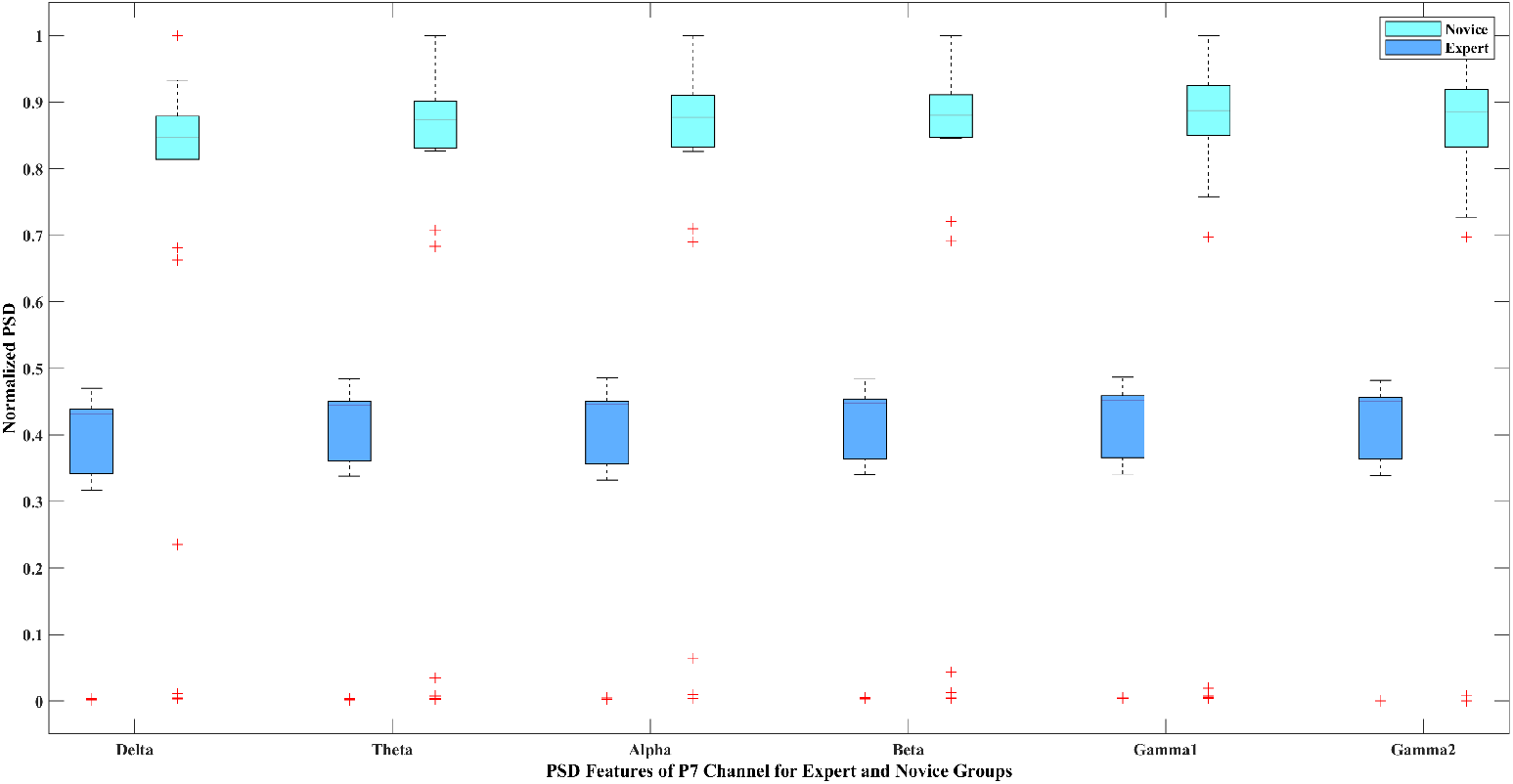
Box-plot representation of the PSD features of *P*7 channel for expert and novice groups.

## Discussion

This study is aimed at classifying the game player’s expertise while optimizing the classification accuracy and reducing the computational overhead. Our results have shown that the proposed scheme achieves an accuracy of 98.04% by utilizing the features of only two EEG channels using the KNN classifier. Moreover, the classification of game player expertise has produced the minimum error of classification. The studies that have performed EEG based game player expertise classification are compared with the proposed framework in Table 2. We also performed leave-one-out cross-validation and leave-one-subject-out CV in addition to 10-fold CV. We thereby evaluated the results from the 10-fold CV in terms of accuracy. It should be noted that since the number of training instances was limited, LOOCV was feasible and gave us insights into the classifier performance. The classification accuracy achieved from LOOCV using the KNN classifier was similar to 10-fold CV. While for LOSO, the classification accuracy dropped but was still significant. The performance is bound to increase as more subjects are added. While we observe a drop in performance, but it must be emphasized, multiple trials were performed to augment the data, and we consider each trial instance as an independent subject when performing 10-fold CV. Moreover, the comparison is performed based on the number of electrodes, subjects, stimulus, classification algorithm, and average classification accuracy achieved.

**Table 2.** A comparison of the current study with related studies in game research.

| Method | Sensors | Subjects | Stimulus | Classifier | Validation | Accuracy (%) |
| --- | --- | --- | --- | --- | --- | --- |
| [40] | 14 | 8 | Guitar Hero | Logistic Regression | 10-fold | 80 |
| [42] | 14 | 10 | Temple Run | NB | 10-fold | 89.89 |
| [41] | 14 | 20 | Temple Run | NB | 10-fold | 88 |
| [43] | 14 | 8 | 3-D modelling | KNN | 5-fold | 95 |
| Proposed | 2 | 10 | Temple Run | KNN | LOSO | 88 |
| Proposed | 2 | 10 | Temple Run | KNN | LOOCV | 98.04 |
| Proposed | 2 | 10 | Temple Run | KNN | 10-fold | 98.04 |

[40] has classified the game player’s expertise using the video game *Guitar Hero* which requires the attention and coordination to play the musical notes on the musical songs of the *rock* genre. The study was conducted on eight participants, out of which four were experts, and four were novices. Participants were asked to play two songs in two modes, i.e., easy and difficult. Delta, theta, alpha, and beta band, along with the two ratios of delta/beta and theta/beta, were used as the features for game player expertise classification. Their results showed that expert players’ cognitive activity is different from that of novices, and the accuracy achieved for game player expertise classification is 80%. The study conducted in [42] has performed the EEG based game player expertise classification using the time-domain features and machine learning approach. The EEG data were acquired from the ten participants while playing *Temple Run* mobile game using the 14-channel Emotiv EPOC headset. Thirteen-time domain features were extracted from each of the fourteen channels resulting in a feature matrix of 50 × 182. The classification was performed using the SVM, NB, and MLP classifiers. Their results showed that the NB classifier best classifies the game player expertise with 89.89% classification accuracy. The study conducted in [41] has performed the game player expertise classification while aiming for the feature-length reduction. The EEG data were acquired using a 14-channel Emotiv EPOC headset from twenty participants for their five trials of gameplay of *Temple Run* mobile game. Thirteen-time domain features were extracted from all fourteen EEG channels. The analysis was performed using all fourteen and four EEG channels for the game player expertise classification, and their performance was analyzed carefully. Their results indicated that the two-player groups’ brain activity was different during the period of gameplay. Moreover, their results showed that with the feature matrix of 50 × 182, SVM achieved 86% classification accuracy while the feature matrix of 50 × 52 NB classified the game player expertise with the 88% classification accuracy. The study performed in [43] is not game-based, but it has proposed a framework of expertise classification in the 3D environment using the Normalized Transfer Entropy (NTE) of the EEG features. This study’s classification accuracy is 95% using five significant features extracted from the EEG data of the frontal electrodes. Comparable to the previous studies, our proposed framework of game player expertise classification is based on the frequency domain analysis that utilizes EEG bands as features from the relevant EEG channels obtained by the *Correlation-based feature selection*. The result of the current study is analyzed using a 10-fold cross-validation model. Our proposed framework outperforms the studies of [40–43] and utilizes the features of only two EEG channels, AF3 and P7. We utilized six features from the two EEG channels and achieved a classification accuracy of 98.04% through the KNN classifier.

Despite obtaining the highest accuracy of classification, the results of our analysis should be considered with some limitations to understand the scope and applicability of our research. Nonetheless, a larger sample of data is needed to expand the current findings and develop methodological procedures to classify the expertise of the game player in the various game genre. The shortcomings of the current study should be further validated through experiments utilizing multiple game stimuli. Moreover, the effect of movement, environment, and system parameters can also be incorporated to see the effect on the behavior of expert-novice gameplay. Moreover, while a high accuracy is achieved, such experimental protocols have to be carefully designed for clinical application. In particular, the accuracy of wearable EEG headsets needs careful attention when dealing with more critical application areas.

The findings of the current study will be beneficial for the development of adaptive games that adjust themselves as per the physiological changes occurring in the game player. Furthermore, the brain activity of the different game player groups can be supplemented in the game design (as in the neurofeedback games), which will gauge the player to preserve his interest and engagement. Such kind of adaptive game designs are found to be more engaging and help the game players to achieve the desired skill-competence balance as well as improve the attention and cognition of the game player [22, 28]. In the future, we intend to overcome the limitations of the current study by increasing the sample size and utilizing a diverse set of games. At the same time, the analysis presented here has emphasized the applicability and usability of EEG signals acquired using wearable devices, such as Emotiv EPOC headset, towards evaluating game player expertise. Our experimental results are found to be significant, and hence, in the future, we intend to evaluate various modalities of game player experience.

## Conclusion

This study presents EEG based game player expertise classification using an objective, reliable, and discreet method which is less prone to subjective or biased opinions. Moreover, while working with the EEG signals, it is desirable to reduce the computational overhead. Our experimental results demonstrated that expert and novice game players’ brain activity has noticeable differences, and the expertise of game players is classified with 98.04% accuracy using the KNN classifier. While this is achieved by utilizing features from two EEG channels: AF3 and P7. These findings could be used in the development of DDA based neuro-feedback games, which adapt their game design in correspondence with the changes occurring in the physiological and electrophysiological signals of the game player. Further, digital interfaces, including video and mobile games, can be made more engaging and interactive by incorporating the player’s expertise and experience as a part of the game design. Thus, our results emphasize the efficacy of game players’ cognitive and behavioral aspects to design games that keep the player engaged and motivated during gameplay. In the future, we intend to predict the game score of a game player using EEG signals and regression models. Furthermore, we shall collect more data from subjects engaging with a diverse set of mobile games for an in-depth analysis of game users. Moreover, the engagement behavior of the game player will be investigated using EEG based engagement indices.

## Acknowledgments

This research was funded by DSR, King Abdul Aziz University, Jeddah, Saudia Arabia. Authors are thankful to DSR for the technical and financial support.

## References

1. Association ES, et al.. Essential facts about the computer and video game industry; 2018.

2. global games market N. Newzoo global games market report, trends, insights, and projections towards 2020; 2017.

3. Łupkowski P, Krajewska V. Immersion level and bot player identification in a multiplayer online game: The World of Warships case study. Czasopismo ludologiczne Polskiego Towarzystwa Badania Gier. 2018; p. 155.

4. Huynh S, Kim S, Ko J, Balan RK, Lee Y. EngageMon: Multi-Modal Engagement Sensing for Mobile Games. Proceedings of the ACM on Interactive, Mobile, Wearable and Ubiquitous Technologies. 2018;2(1):1–27.

5. Gerling KM, Klauser M, Niesenhaus J. Measuring the impact of game controllers on player experience in FPS games. In: Proceedings of the 15th International Academic MindTrek Conference: Envisioning Future Media Environments. ACM; 2011. p. 83–86.

6. Boyle EA, Connolly TM, Hainey T, Boyle JM. Engagement in digital entertainment games: A systematic review. Computers in human behavior. 2012;28(3):771–780.

7. Jennett C, Cox AL, Cairns P, Dhoparee S, Epps A, Tijs T, et al. Measuring and defining the experience of immersion in games. International journal of human-computer studies. 2008;66(9):641–661.

8. IJsselsteijn W, De Kort Y, Poels K. The game experience questionnaire. Eindhoven: Technische Universiteit Eindhoven. 2013;.

9. Brockmyer JH, Fox CM, Curtiss KA, McBroom E, Burkhart KM, Pidruzny JN. The development of the Game Engagement Questionnaire: A measure of engagement in video game-playing. Journal of Experimental Social Psychology. 2009;45(4):624–634.

10. Ryan RM, Rigby CS, Przybylski A. The motivational pull of video games: A self-determination theory approach. Motivation and emotion. 2006;30(4):344–360.

11. Denisova A, Nordin AI, Cairns P. The convergence of player experience questionnaires. In: Proceedings of the 2016 Annual Symposium on Computer-Human Interaction in Play. ACM; 2016. p. 33–37.

12. Diva SZ, Prorna RA, Rahman II, Islam AB, Islam MN. Applying brain-computer interface technology for evaluation of user experience in playing games. In: 2019 International Conference on Electrical, Computer and Communication Engineering (ECCE). IEEE; 2019. p. 1–6.

13. Maby E, Perrin M, Bertrand O, Sanchez G, Mattout J. BCI could make old two-player games even more fun: a proof of concept with “Connect Four”. Advances in Human-Computer Interaction. 2012;2012.

14. Arellano DG, Tokarchuk L, Gunes H. Measuring affective, physiological and behavioural differences in solo, competitive and collaborative games. In: International conference on intelligent technologies for interactive entertainment. Springer; 2016. p. 184–193.

15. Kneer J, Elson M, Knapp F. Fight fire with rainbows: The effects of displayed violence, difficulty, and performance in digital games on affect, aggression, and physiological arousal. Computers in Human Behavior. 2016;54:142–148.

16. Nacke LE, Grimshaw MN, Lindley CA. More than a feeling: Measurement of sonic user experience and psychophysiology in a first-person shooter game. Interacting with computers. 2010;22(5):336–343.

17. Nacke LE. An introduction to physiological player metrics for evaluating games. In: Game analytics. Springer; 2013. p. 585–619.

18. Kivikangas JM, Chanel G, Cowley B, Ekman I, Salminen M, Järvelä S, et al. A review of the use of psychophysiological methods in game research. journal of gaming & virtual worlds. 2011;3(3):181–199.

19. Khairuddin HR, Malik AS, Mumtaz W, Kamel N, Xia L. Analysis of EEG signals regularity in adults during video game play in 2D and 3D. In: 2013 35th Annual International Conference of the IEEE Engineering in Medicine and Biology Society (EMBC). IEEE; 2013. p. 2064–2067.

20. Nacke LE, Stellmach S, Lindley CA. Electroencephalographic assessment of player experience: A pilot study in affective ludology. Simulation & Gaming. 2011;42(5):632–655.

21. Abhishek AM, Suma H. Stress analysis of a computer game player using electroencephalogram. In: International Conference on Circuits, Communication, Control and Computing. IEEE; 2014. p. 25–28.

22. Chanel G, Rebetez C, Bétrancourt M, Pun T. Emotion assessment from physiological signals for adaptation of game difficulty. IEEE Transactions on Systems, Man, and Cybernetics-Part A: Systems and Humans. 2011;41(6):1052–1063.

23. Tanaka T, Arvaneh M. Signal processing and machine learning for brain-machine interfaces. Institution of Engineering and Technology; 2018.

24. Nijholt A, Bos DPO, Reuderink B. Turning shortcomings into challenges: Brain–computer interfaces for games. Entertainment computing. 2009;1(2):85–94.

25. Schoneveld EA, Malmberg M, Lichtwarck-Aschoff A, Verheijen GP, Engels RC, Granic I. A neurofeedback video game (MindLight) to prevent anxiety in children: A randomized controlled trial. Computers in Human Behavior. 2016;63:321–333.

26. Wang Q, Sourina O, Nguyen MK. Fractal dimension based neurofeedback in serious games. The Visual Computer. 2011;27(4):299–309.

27. Sekhavat YA. Battle of minds: a new interaction approach in BCI games through competitive reinforcement. Multimedia Tools and Applications. 2020;79(5):3449–3464.

28. Thomas KP, Vinod A, Guan C. Enhancement of attention and cognitive skills using EEG based neurofeedback game. In: 2013 6th International IEEE/EMBS Conference on Neural Engineering (NER). IEEE; 2013. p. 21–24.

29. Derbali L, Frasson C. Prediction of players motivational states using electrophysiological measures during serious game play. In: 2010 10th IEEE International Conference on Advanced Learning Technologies. IEEE; 2010. p. 498–502.

30. Bakaoukas AG, Coada F, Liarokapis F. Examining brain activity while playing computer games. Journal on Multimodal User Interfaces. 2016;10(1):13–29.

31. Mishra J, Zinni M, Bavelier D, Hillyard SA. Neural basis of superior performance of action videogame players in an attention-demanding task. Journal of Neuroscience. 2011;31(3):992–998.

32. Parsons TD, McMahan T, Parberry I. Neurogaming-based classification of player experience using consumer-grade electroencephalography. IEEE Transactions on Affective Computing. 2015;.

33. Xu J, Zhong B. Review on portable EEG technology in educational research. Computers in Human Behavior. 2018;81:340–349.

34. Vasiljevic GAM, de Miranda LC. Brain–computer interface games based on consumer-grade EEG Devices: A systematic literature review. International Journal of Human–Computer Interaction. 2020;36(2):105–142.

35. Loh CS, Sheng Y, Li IH. Predicting expert–novice performance as serious games analytics with objective-oriented and navigational action sequences. Computers in Human Behavior. 2015;49:147–155.

36. Berta R, Bellotti F, De Gloria A, Pranantha D, Schatten C. Electroencephalogram and physiological signal analysis for assessing flow in games. IEEE Transactions on Computational Intelligence and AI in Games. 2013;5(2):164–175.

37. Rebetez C, Betrancourt M. Video game research in cognitive and educational sciences. Cognition, Brain, Behavior. 2007;11(1):131–142.

38. Jones C, Scholes L, Johnson D, Katsikitis M, Carras MC. Gaming well: links between videogames and flourishing mental health. Frontiers in psychology. 2014;5:260.

39. Stanhope JL, Owens C, Elliott LJ. Stress Reduction: Casual Gaming versus Guided Relaxation. 2016;.

40. Lujan-Moreno GA, Atkinson R, Runger G, Gonzalez-Sanchez J, Chavez-Echeagaray ME. Classification of video game players using EEG and logistic regression with ridge estimator. In: Workshop on Utilizing EEG Input in Intelligent Tutoring Systems (ITS2014 WSEEG); 2014. p. 21.

41. Anwar S, Saeed S, Majid M, Usman S, Mehmood C, Liu W. A game player expertise level classification system using electroencephalography (EEG). Applied Sciences. 2018;8(1):18.

42. Anwar SM, Saeed SMU, Majid M. Classification of Expert-Novice Level of Mobile Game Players Using Electroencephalography. In: 2016 International Conference on Frontiers of Information Technology (FIT). IEEE; 2016. p. 315–318.

43. Baig MZ, Kavakli M. Expertise Classification using Functional Brain Networks and Normalized Transfer Entropy of EEG in Design Applications. In: Proceedings of the 2019 11th International Conference on Computer and Automation Engineering. ACM; 2019. p. 41–46.

44. Hall MA. Correlation-based feature selection for machine learning. 1999;.

45. Welch P. The use of fast Fourier transform for the estimation of power spectra: a method based on time averaging over short, modified periodograms. IEEE Transactions on audio and electroacoustics. 1967;15(2):70–73.

46. Gunn SR, et al. Support vector machines for classification and regression. ISIS technical report. 1998;14(1):5–16.

47. John GH, Langley P. Estimating continuous distributions in Bayesian classifiers. In: Proceedings of the Eleventh conference on Uncertainty in artificial intelligence. Morgan Kaufmann Publishers Inc.; 1995. p. 338–345.

48. Lin YP, Wang CH, Jung TP, Wu TL, Jeng SK, Duann JR, et al. EEG-based emotion recognition in music listening. IEEE Transactions on Biomedical Engineering. 2010;57(7):1798–1806.

49. Murugappan M. Human emotion classification using wavelet transform and KNN. In: 2011 International Conference on Pattern Analysis and Intelligence Robotics. vol. 1. IEEE; 2011. p. 148–153.

50. Thenmozhi M. Forecasting stock index returns using neural networks. Delhi Business Review. 2006;7(2):59–69.

51. Mallapragada C. Classification of EEG signals of user states in gaming using machine learning. 2018;.

52. Chayer C, Freedman M. Frontal lobe functions. Current neurology and neuroscience reports. 2001;1(6):547–552.

